# Cell motion as a stochastic process controlled by focal contacts dynamics

**DOI:** 10.1101/750331

**Authors:** Simon Lo Vecchio, Raghavan Thiagarajan, David Caballero, Vincent Vigon, Laurent Navoret, Raphaël Voituriez, Daniel Riveline

**Affiliations:** Laboratory of Cell Physics ISIS/IGBMC, CNRS and University of Strasbourg, Strasbourg, France; Institut de Génétique et de Biologie Moléculaire et Cellulaire, Illkirch, France; Centre National de la Recherche Scientifique, UMR7104, Illkirch, France; Institut National de la Santé et de la Recherche Médicale, U964, Illkirch, France; Université de Strasbourg, Illkirch, France; Institut de Recherche Mathematique Avancee, UMR 7501, Universite de Strasbourg et CNRS; Laboratoire de Physique Théorique de la Matière Condensée, UMR 7600 CNRS/Sorbonne Université, 4 Place Jussieu, 75255 Cedex Paris, France; Laboratoire Jean Perrin, UMR 8237 CNRS/Sorbonne Université, 4 Place Jussieu, 75255 Cedex Paris, France

## Abstract

Directed cell motion is essential in physiological and pathological processes such as morphogenesis, wound healing and cancer spreading. Chemotaxis has often been proposed as the driving mechanism, even though evidence of long-range gradients is often lacking in vivo. By patterning adhesive regions in space, we control cell shape and the associated potential to move along one direction in another mode of migration coined ratchetaxis. We report that focal contacts distributions collectively dictate cell directionality, and bias is non-linearly increased by gap distance between adhesive regions. Focal contact dynamics on micro-patterns allow to integrate these phenomena in a consistent model where each focal contact can be translated into a force with known amplitude and direction, leading to quantitative predictions for cell motion in every condition. Altogether, our study shows how local and minutes timescale dynamics of focal adhesions and their distribution lead to long term cellular motion with simple geometric rules.

## INTRODUCTION

The ability of cells to perform directed motion is critical to the proper function of many physiological, pathological or developmental processes (1-4). Bias in cell migration have been widely studied *in vitro* and it has been shown that soluble gradients of specific chemicals could trigger directionality by regulating signalling pathways and reorganizing the acto-myosin cytoskeleton (5,6). This phenomenon, known as *chemotaxis*, remains the widespread way to explain cell directionality *in vivo* (7-9). However, these gradients are rarely demonstrated or quantified *in vivo*. In addition, if they are reported, they are rarely shown to be *causal* for cell directions. How cells can sustain directions *in vivo* remains therefore unclear.

Previous works showed *in vitro*, as a proof of a principle, that local asymmetries in cellular environment can bias cell directionality in another mode of migration named *ratchetaxis* (10-17). This type of behavior can lead to long-term directionality and stresses the importance of stochastic probing associated to cell protrusions, as well as the role of environment topology. Using *ratchetaxis* as our framework, we show that cell directionality is primarily determined by the collective behavior of its adhesion sites – namely focal contacts – and can be tuned and predicted using simple geometric rules describing the surrounding environment. We also define a critical nucleation area, responsible of the observed bias: below this area, protrusions retract and the motion is prohibited, and above this area, new focal adhesions nucleate in a “wave-like” manner and cell migrates. This framework is supported by a theoretical model integrating accessible parameters and fitting the rectification plots extracted from experiments.

## RESULTS

Cells locally elongate protrusions, which eventually lead to adhesion and its subsequent force generation and motion as a whole. To test this interplay between cell protrusion dynamics and cell motion, we designed an assay where an array of triangular adhesive motifs over a range of gap distances were fabricated using standard microfabrication techniques (see Methods). Triangular rhodamine-labelled fibronectin (FN) motifs were patterned to guide adhesion and morphodynamics of NIH3T3 cells (see Figure 1a). Non-printed areas were passivated with poly(L-lysine) grafted poly(ethylene glycol) (PLL-g-PEG), so that cells mainly adhere on fibronectin motifs and have their shapes imposed during the spreading phase. We studied the long-term migration of NIH3T3 fibroblasts along the micro-patterns for 48 h. Cells were able to move from one motif to the neighboring motifs, and eventually exhibit persistent motion towards one particular direction (see Figure 1 b-e and Movies S1-S4). Interestingly, cells migrated faster on separated motifs than on connected ones (see Figure 1h), consistent with the fact that friction is increased with larger adhesive areas (18).

**Figure 1:**
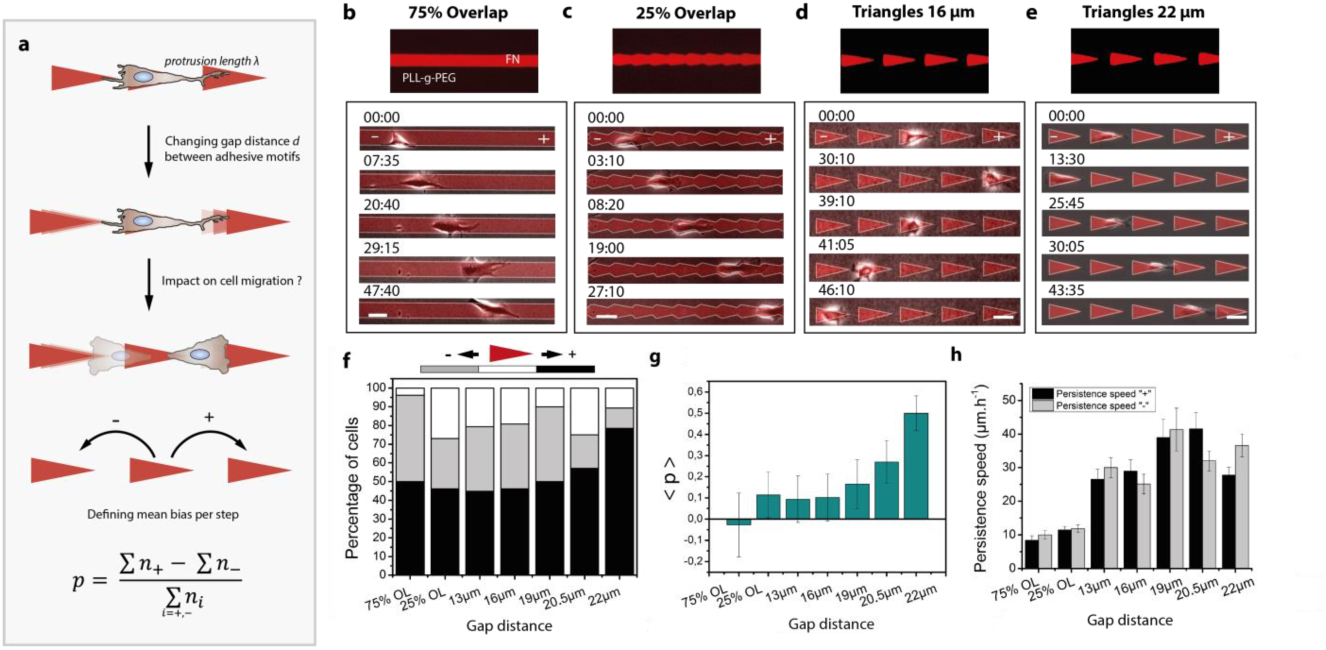
Cell motion depends on gap distance. (a) Scheme showing the experimental setup where a cell is seeded on a microcontact printed array of asymmetric FN triangles. The gap distance is systematically increased, which determines the final direction of migration. Quantitatively, *p* describes the bias in directionality taking into account all steps made during the motion. (b-e) Cells are able to move from one motif to another on separated triangles, and crawl on connected ones. Time in hh:mm, scale bars: 50 µm. (f) Final bias in percentage as a function of gap distance. (g) Average *p* given by Eq. 1 for each condition. (h) Persistence speed in µm.h^-1^. Data set: 25 < N_cells_ < 30 and N_biological_ > 5 for 75% OL, 25% OL, 13µm, 16µm, 19µm and 22µm. N_cells_ = 16 for 20.5 µm. Data is shown as Mean ± S.E.

For each cell moving along a lattice unit (defined as the distance between the centers of adjacent triangles), we defined a final *bias*, which is determined by the position of the cell at the end of its trajectory relative to its original location after 2 days of migration: the final bias for an individual cell was set as “+” if it migrated towards the tip of the motif, and “–” if it migrated towards base of the motif (see Figure 1f). Interestingly, while the bias was not significant for triangles separated by 13 µm, 16 µm and 19 µm gap distances, we found that the average bias increased and reached close to complete *rectification* on triangles separated by gaps of 22 µm: almost 80% of the cells moved, on average, towards the “+” direction in this condition (Figure 1f). Beyond this value, the distance exceeded the maximum extension of protrusions. We tested large gap distances (d = 45 µm), and as expected, no migration was observed (see Movie S5); cells were unable to extend protrusions and find a ‘docking’ site. These results show the sharp increase of rectification as a function of gap distance.

We next sought to quantify the complete trajectory of cells for each experimental condition where cells can move back and forth during 2 days. We computed the bias *p* per lattice unit defined as:

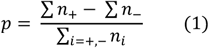

where *n*_*i*_ corresponds to the number of steps performed in the *i* direction. This readout reports how cells behave for each gap distance independently of the total duration of a trajectory. We averaged over trajectories, and the resulting < *p* > confirms the behavior observed for the total bias (see Figure 1g). For 22 µm, the total bias is maximal, and the mean bias per step is also sharply increased and reaches 0.48 (± 0.08). Distributions of *p* are also informative: the distribution obtained for the line of fibronectin (75% OL) corresponds to a stochastic bistable behavior with two peaks around p = 1 and p = -1 (see Figure S1) and no preferred direction. This feature progressively vanishes and bistability is transformed into a monostable distribution with one single peak located around p = 1 for 22 µm. Altogether, these results suggest that cell micro-environment architecture and simple geometrical features are sufficient to tune bistability into an almost deterministic cell directionality, and importantly, in the absence of chemical gradients.

Focal adhesions (FA) are protein complexes, connected to the actomyosin meshwork, acting as an adhesive anchor and as a local force transmitter (19-20). Considering the monostable condition (22 µm) being the most simple and efficient configuration, we next asked whether FAs dynamics and distribution could be sufficient to explain rectification. Using NIH3T3 fibroblasts stably expressing GFP focal-adhesion vasodilator-stimulated phosphoprotein (VASP, see Methods and Figure S2), we recorded the dynamics of adhesion sites during motion on the micro-patterned fibronectin motifs (see Movie S6). We observed that all protrusions, as soon as they touch fibronectin regions, generate FAs (see Figure 2 and Movie S6). When the first row of FAs nucleates on the neighbouring adhesive pattern, new contacts appear along the motif in a wave-like manner (see Figure S3 and Movie S7). This indicates a *collective* behaviour. This phenomenon on adhesive motifs is distinct from the protrusions growing on PLL-g-PEG. In this latter case, protrusion growth stops and retracts after nucleation of FAs. This suggests that FAs dynamics could be a relevant readout for cell motion.

**Figure 2:**
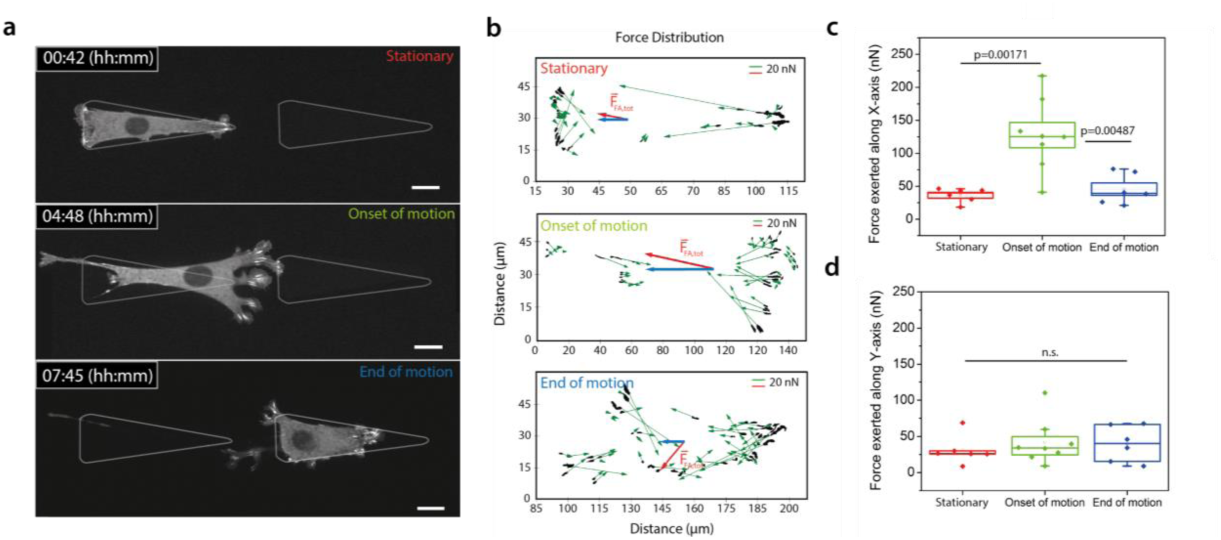
Force inference based on focal contacts distributions correlates with motion. (a) Time-lapse images of an NIH3T3 cell stably expressing VASP-GFP. Three time points are extracted for analysis: (i) during the stationary phase, (ii) at the onset of motion, and (iii) at the end of motion. (b) Force vectors map. Each focal contact is linked to a force vector oriented in the direction of adhesion zone (in green). Its amplitude is set according to the relationship 1 µm^2^ corresponds to 5.5 nN. The total force vector is displayed in red, its x component in blue; (c-d) Force exerted along x-axis and y-axis during the three time points. Statistical tests comparing distributions are done with a one-way ANOVA. N_biological_repeat_ = 4, N_cells_ = 8. Scale bar = 15µm.

We next tested this hypothesis quantitatively. It was shown that focal adhesions act as mechanosensors and can apply a force per unit area of 5.5 nN.µm-2 (19-21), which is conserved across cell types and substrates (20,21). We used this relationship to test whether the force inferred from focal contacts could be predictive of motion. While the global balance of forces applied on the cell is verified at all times (22), the spatial distribution of forces is informative of local dynamics and global cell motion. We mapped force vectors at three different relevant time-points (see Figure 2). Each focal adhesion mediates a force *f*_*i*_ exerted by the cell on the substrate along the long axis of the focal contacts, and its orientation points towards the interior of the cell. Its magnitude is deduced from the measured area of the FA by assuming a constant stress of 5.5 nN.µm-2 (Figure 2b). From these vectors, we extracted the total force associated with each focal contact, its amplitude and its direction along x and y axis for different cells (Figure 2c,d). Cells apply a large total force of 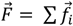 and is about 130 nN along the x-axis during motion, 3-fold larger than prior and after motion. To satisfy global force balance, a resistive friction force has to be taken into account. Assuming viscous friction, the corresponding friction coefficient can be estimated as ∼ 25 N.s.m-1, also consistent with reported values of friction of ∼ 30 N.s.m-1 (18). This suggests that cell motion can be captured with only focal contacts dynamics translated into local forces and could be explained quantitatively by analysis of distributions of focal contacts right and left. Since every protrusion generates focal adhesions as soon as it touches the patterns (see Figure 3a), distributions of contacts, and therefore cell direction, are expected to be critically governed by adhesive area accessible to protrusions. This was quantified by a simple phenomenological model that expresses explicitly the mean bias per step as a function of adhesive area accessible to protrusions on each side. Assuming a linear dependence of probabilities of formation of a focal adhesion with the adhesive area probed by protrusions made it possible to derive a simple expression of <*p*> involving only measurable geometric parameters ; this however yielded a poor agreement with the plots of <*p*> as a function of gap distance (see Figure 3b and Theory). This indicates that collective effects in focal adhesion formation, as observed (see above) must be taken into account. To do so, we introduced the nucleation area A_0_ (see Theory), a phenomenological parameter representing the typical area that cells need to engage in a neighboring motif to efficiently contribute to motion. For A< A_0_, detachment of the new adhesive zone occurs. Above this area, the nucleating zone is stable and a wave of new contacts is generated. The resulting prediction of < *p* > yielded an excellent agreement with experimental values (see Figure 3b), where notably the single fitting parameter A_0_ was involved.

**Figure 3:**
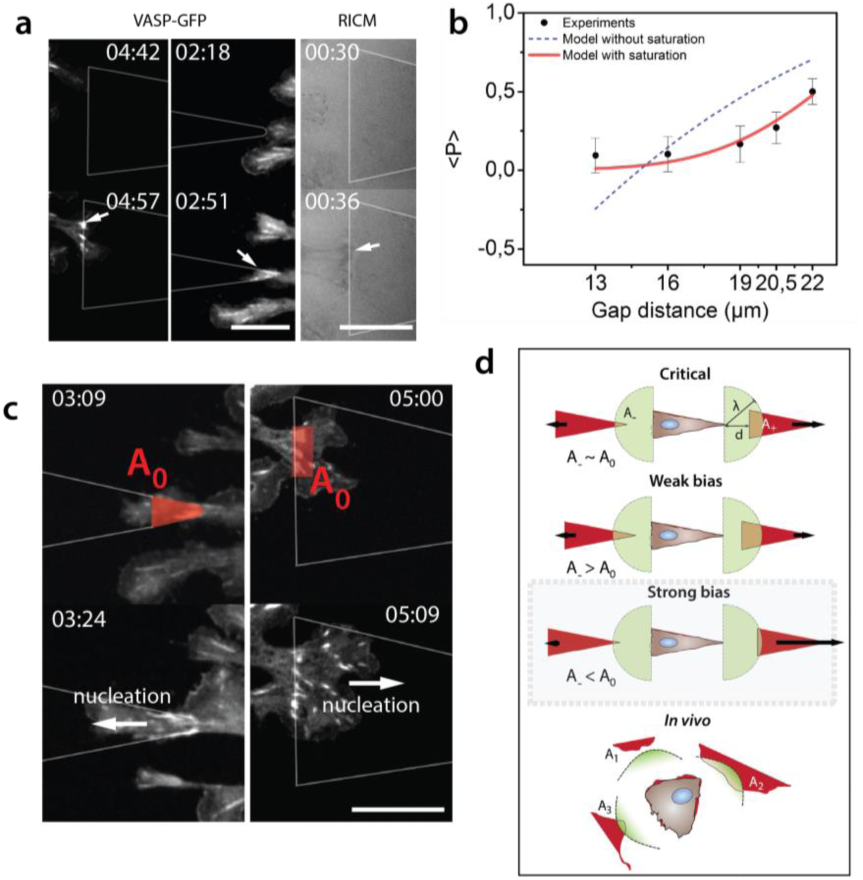
The source of the bias. (a) Protrusions touching an adhesive motif imaged with fluorescence microscopy and RICM. Focal adhesions are nucleated right after contact with motifs. Arrows indicate FA on patterns. (b) Comparisons between model and experiments. The blue fit assumes mechanisms with areas probed by cells, whereas the red fit includes the nucleation area A_0_. Fit corresponds to A_0_=13.5 µm^2^ and λ= 28 µm. (c) Experimental description of A_0_. From this region, subsequent rows of focal adhesions are nucleated. Arrows indicate direction of nucleation. (d) Graphical description of A_0_ and its influence on bias. Beyond A_0_, the bias is weak. If one side gets close or below A_0_, the bias in the opposite direction becomes strong. Time in hh:mm, scale bars = 15µm

We turned back to experiments of focal contact dynamics: the nucleating area A_0_ obtained from the fit of 13.5 µm^2^ indeed matched the nucleation area in experiments (see Figure 3c). This suggests that a nucleation zone is needed to sustain proper anchor on neighboring motifs, consistent with the notion that a collection of focal contacts is required to perform firm cellular anchors on neighboring motifs. Importantly, the model involves only 2 cellular parameters that can be directly measured, and no *ad hoc* adjustable parameter was involved to recapitulate the trend observed in experiments. This supports the relevance of our hypothesis. If the area available on both sides is greater than A_0_, the cell will be able to create stable anchoring zones sufficient to nucleate the subsequent rows of FAs. Therefore, the bias will be poor even if A_+_ ≪ A_-_ (13 µm to 19 µm conditions, see Figure 3d). On the other hand, if one side goes close or below A_0_, the bias becomes stronger in the opposite direction. This kind of coupling between geometry and cellular features could be relevant *in vivo* where distributions of adhesive regions are not homogeneous (Figure 3d and (23)).

## DISCUSSION

Overall, our study shows that cell migration can be tuned and captured by simple geometric rules associated with cellular features. So far, focal contacts have been studied mainly individually, and their integrated action was often overlooked. Our results illustrate how collective behavior of focal adhesions can mediate cell motion and establishes new rules for cell direction from the coupled dynamics of focal contacts and the cellular environment.

Ratchetaxis relevance could be probed in the future *in vivo*, along lines recently suggested (23). To test this framework, anisotropy in adhesion sites need to be tracked. In addition, protrusions and cell contacts dynamics have to be acquired and correlated over time quantitatively, together with long term cell motion. These read-outs are now accessible *in vivo* for a variety of model systems, but their simultaneous observations and correlations are not performed yet to our knowledge. New experiments will need to be conducted to establish this potential source of rectification for cell migration *in vivo*.

In addition to its novel way of operating, ratchetaxis encodes cellular mechanisms which convey new ways of perceiving cell motion with readouts extracted from biological physics. Cell *polarity* is acquired in the first steps through interactions with environment, and then a *persistence in polarity* is kept over time by the cell. Eventually, this translates into the mean direction taken by cells through collective effect of focal contacts. These parameters, polarity, persistence of polarity, and friction quantified through focal contacts, can be integrated in predictive models, as we reported in this study. Also, these steps and associated parameters are all stochastic, and their distributions are informative to test hypothesis. Altogether, ratchetaxis could also serve as a key phenomenon to test physical models for living matter.

## ACKNOWLEDGMENTS

We thank O. Pertz, P. Rossolilo and A. Dupont, the Imaging Platform of IGBMC, the Riveline Lab. for constructs and discussions. This study with the reference ANR-10-LABX-0030-INRT has been also supported by a French state fund through the Agence Nationale de la Recherche under the frame programme Investissements d’Avenir labelled ANR-10-IDEX-0002-02. SLV is supported by the University of Strasbourg. R.T. was an IGBMC International PhD Programme fellow supported by LabEx INRT funds. D.R. acknowledges support from ciFRC Strasbourg, the University of Strasbourg, CNRS and Foundation Cino del Duca.

## Conflicts of interest

none.

## Materials and Methods

### Micro-contact printing

Fibronectin asymmetric motifs were patterned using standard microcontact printing protocol (24-25). Briefly, polydimethylsiloxane (PDMS, 1:10 w/w cross-linker:pre-polymer) (Sylgard 184 kit, Dow Corning, cat. DC184-1.1) stamps were replicated from SU-8 molds fabricated by standard UV-photolithography. After cleaning in ethanol 70%, stamps were rendered hydrophilic by oxygen plasma activation (Diener Electronic, cat. ZeptoB) for 30 s. Then, stamps were incubated for 1h with a 10 µg/mL rhodamine-labelled fibronectin solution (Cytoskeleton, cat. FNR01-A) and dried under a stream of nitrogen at room temperature for about 5 min. Next, stamps were placed in contact on top of a glass coverslip functionalized by vapour phase for 1 h with 3-(mercapto)propyltrimethoxysilane (FluroChem). A 50 g weight was placed on the top of the stamp during 30 min to ensure a constant and isotropic pressure during stamping. After release, the coverslip was cleaned in PBS 1x. The non-patterned areas were passivated with a 0.1 mg/mL solution of PLL-g-PEG (in 1 mM HEPES pH 7.4, SuSoS AG, cat. SZ33-15) for 20 min at room temperature. Finally, the patterned glass coverslips were again rinsed in PBS 1x prior cell deposition.

### Cell Culture

NIH3T3 mouse fibroblasts (ATCC) were cultured in high glucose Dulbecco’s Modified Eagle’s Medium (DMEM) (Fisher Scientific, cat. 11574486) supplemented with 10% Bovine Calf Serum (BCS, Sigma, cat. 12133C) and 1% Penicillin-Streptamycin. Cells were detached from the Petri dish with Trypsin-0.25% EDTA (Fisher Scientific, cat. 11570626), centrifuged, and re-suspended in DMEM 10% BCS. About 2000 cells were seeded on micro-patterns and the non-adhering cells were washed out after 30 min of incubation at 37°C and 5% CO_2_. Medium was replaced by L-15 (Leibovitz Medium, Fisher Scientific, cat. 11540556) supplemented with 1% of BCS in order to reduce cell division during experiments.

Focal adhesions, named also focal contacts, were monitored by fluorescence microscopy. NIH3T3 mouse fibroblasts stably expressing a VASP-GFP vector were kindly provided by Olivier Pertz (Institute of Cell Biology, Bern University, Switzerland) (26). The construct is expressed in lentivirus derived from pLenti CMV MVS. The viruses were added to a culture of wild-type NIH3T3 and cells were selected after 24 h with 2 µg/mL of puromycin.

We checked that GFP-VASP was a direct and reliable readout of focal adhesions (see Methods). This was confirmed by immuno-staining cells expressing GFP-VASP with paxillin. Cells were fixed with 4% paraformaldehyde for 8 min. Then, they were treated with Triton 0.5X during 3 min. After washout, an anti-paxillin mouse primary antibody (BD transduction, cat. 610051) was used together with an anti-mouse secondary antibody grafted with Cy3 (Jackson Immuno Research, cat. 115-165-146). We observed co-localisation providing us good evidence that GFP-VASP reveals focal contacts – see also results with RICM below.

### Time-Lapse microscopy

Long-term phase-contrast images (48 h) were acquired using a low magnification objective (4x, N.A. 0.25) with an image acquisition frequency of 1 image each 5 min. Micro-patterns were visualised by means of a standard epifluorescence lamp (FluoArc Hg Lamp) coupled with a rhodamine fluorescence filter. A thin layer of mineral oil (Sigma, M5904) was deposited on top of the cell culture media in order to prevent medium evaporation. In Figure 1, cells were imaged using a 10x objective (N.A. 0.4). Focal contacts dynamics were monitored every 3 min with a Nikon Spinning-Disk confocal microscope with a 60x oil objective (N.A. 1.4) using the Perfect Focus System. These settings provided highly resolved images with a low phototoxicity for long-term experiments (15 h). To test whether VASP-GFP protein revealed focal contacts localisations, we used Reflection Interference Contrast Microscopy (RICM) as described in Curtis et al. (27). We used a homogeneous white light source (FluoArc Hg Lamp) through a 50% dichroic and imaged interferences on the surface (see Figure S2b).

### Image processing

Tracking of the cells (nucleus) was performed with ImageJ using Multiple Tracking plug-in. The generated coordinates were used to compute cell trajectories. Plots derived from data as well as statistical tests were done using OriginLab Pro (OriginLab). Force analysis made on focal contacts was performed with a custom Python code computing areas of adhesion zones, their orientations and barycenters, force vectors for each of them and total force vectors taking the rule 1 µm^2^ corresponds to 5.5 nN (20-21). Fits with experimental data were done through OriginLab Pro.

### Theory

At low Reynolds number, force balance for each cell can be written (22):

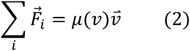

where 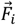 denotes the force exerted by the substrate on the cell through a given focal adhesion *i*, and *µ*, which can be non-linear, effectively accounts for all friction forces resulting from the interaction of cell body with substrate. Importantly, the direction of motion is here fully determined by the knowledge of 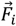, which can be determined quantitatively from direct imaging of focal adhesions, as argued above, and be used as a tool to predict the direction of motion.

Projecting Eq. 1 along the ratchet axis and averaging over time yields the following relation between the bias per step and mean number of focal adhesions *n*_+_ (*resp*. *n*_−_) in the + (resp. -) direction: *p* ∝ (*n*_+_ − *n*_−_)/(*n*_+_ + *n*_−_). Introducing now the purely geometric quantities A_+_ and A_-_, defined as the area accessible to protrusions and FAs in each direction, we assume that *n*_+,−_ ∝ *nA*_+,−_ (other functional form, provided that n vanishes with A, would yield qualitatively similar results). This yields finally to:

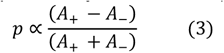

which elucidates the dependence of the bias on geometry. Interestingly, the estimated ratio 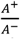 is always greater than 1 for protrusions lengths compatible with NIH3T3 cells and shows a monotonic behavior coupled to a strong divergence when the gap distance is close to the mean protrusion length λ (see Figure S4). When λ is large enough (i.e. close to the size of a pattern), the ratio of probed areas goes back to 1.

When we include the nucleation area A_0_, we used the following phenomenological expression of p:

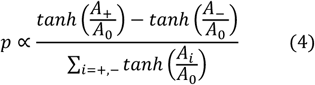

A_0_ can be thought as the critical area above which there is no significant bias because the cell has the opportunity to create a sufficient stable anchoring region to nucleate the subsequent rows of focal contacts on both sides, even if A_+_ ≪ A_-_ (see Figure 3d and Movie S7). The value of A_0_ in our model corresponds to 13.5 µm^2^, and this strikingly corresponds to a single row of focal contacts areas on patterns.

Interestingly, A_i_ is determined with no fitting parameters from the knowledge of the mean protrusion length and the geometry of the patterns. With these minimal number of parameters, we can qualitatively reproduce observations and quantitatively predict a sharp increase of the bias for a gap of 22 mm, as observed in experiments (Figure 3c).

The parameter λ, embedded into the computations of A_i_, corresponds to the typical length of protrusions (essentially between 20 µm and 35 µm in our experiments for NIH 3T3). To fit our model with experiments, we used λ = 28 µm. Therefore, fitting parameters encode actual cellular quantities, and no *ad hoc* adjustable parameters or global pre-factors are added to fit the measures extracted from experiments. This supports the relevance of our hypothesis.

