## Supplementary figures for "Cell motion as a stochastic process controlled by focal contacts dynamics"

**Movie description**

**Movie S1:** NIH3T3 fibroblast migrating on a fibronectin line (75% OL). hh:mm, scale bar = 50µm

**Movie S2:** NIH3T3 fibroblast migrating on connected triangles (25% OL). hh:mm, scale bar = 50µm

**Movie S3:** NIH3T3 fibroblast migrating on fibronectin triangles separated by 16µm gaps. hh:mm, scale bar = 50µm

**Movie S4:** NIH3T3 fibroblast migrating on fibronectin triangles separated by 22µm gaps. hh:mm

**Movie S5:** NIH3T3 fibroblast fluctuating within a single motif and unable to migrate to the neighboring motifs (45µm). hh:mm

**Movie S6:** Dynamics of focal contacts (VASP-GFP). Cell migrates on fibronectin triangles separated by 22µm gaps. hh:mm, scale bar = 15µm

**Movie S7:** Nucleations of rows of focal adhesions in a wave-like manner

**Supplementary figures**

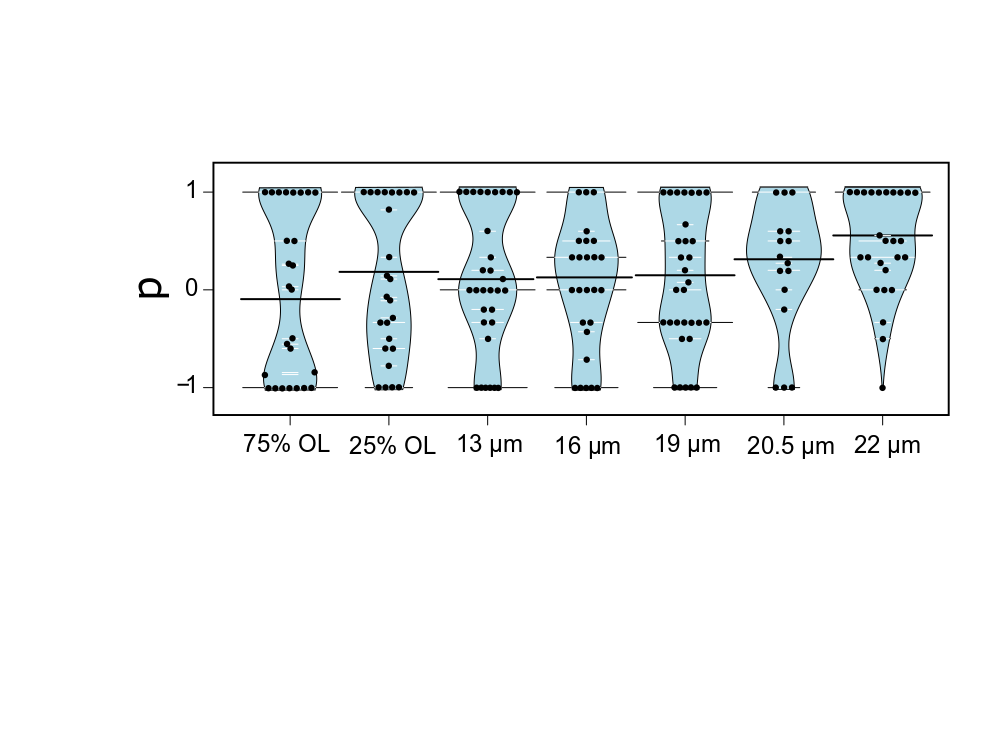

**Figure S1:** Distribution of *p* for all experimental conditions.

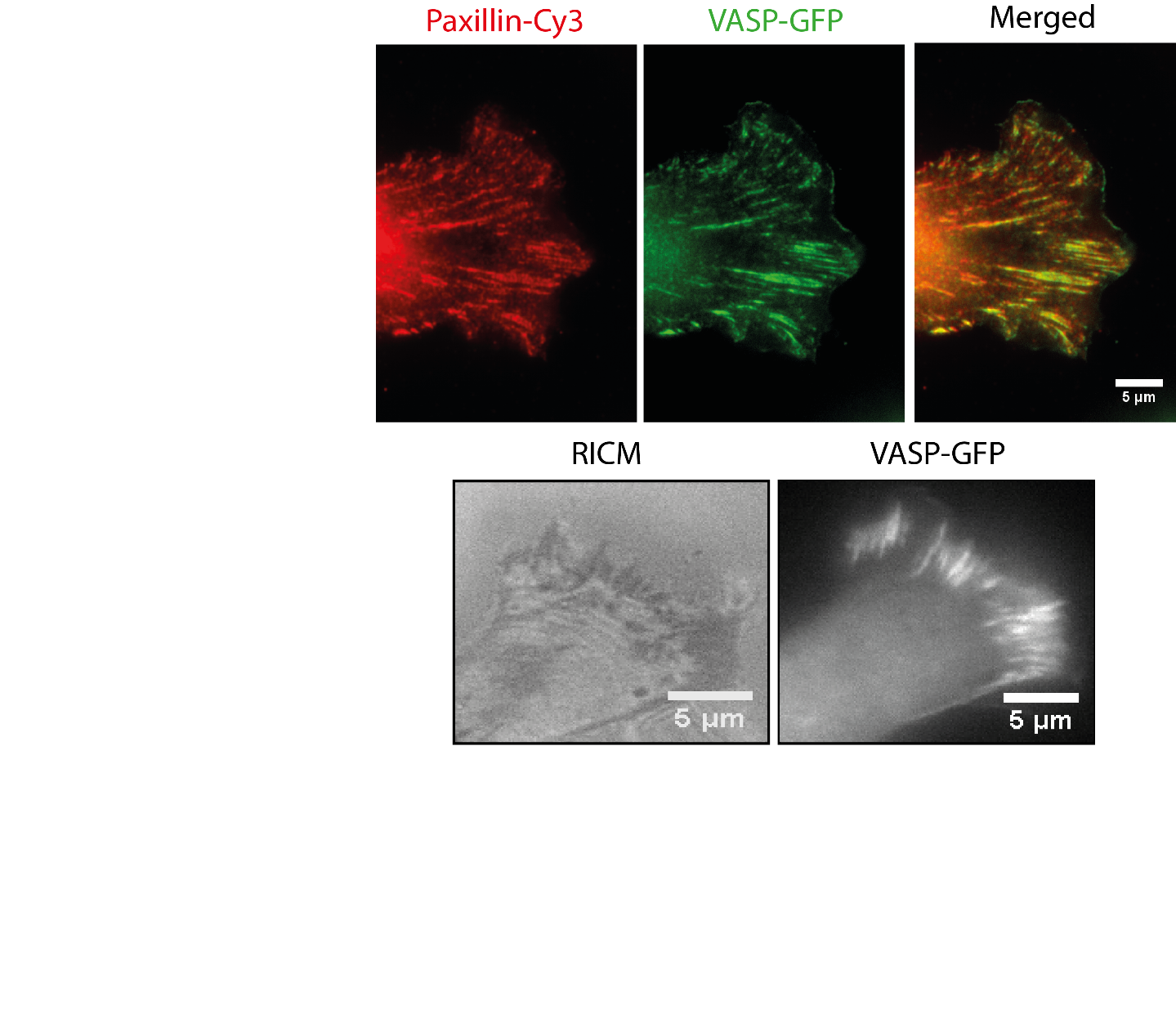

**b**

**a**

**Figure S2: VASP-GFP is a reliable reporter for focal contacts dynamics.** (a) In immunofluorescence microscopy, VASP-GFP and paxillin co-localize. (b) Cells imaged with RICM exhibit the same focal contacts distribution and VASP-GFP co-localize with interferences zones (in dark) within the same cell.

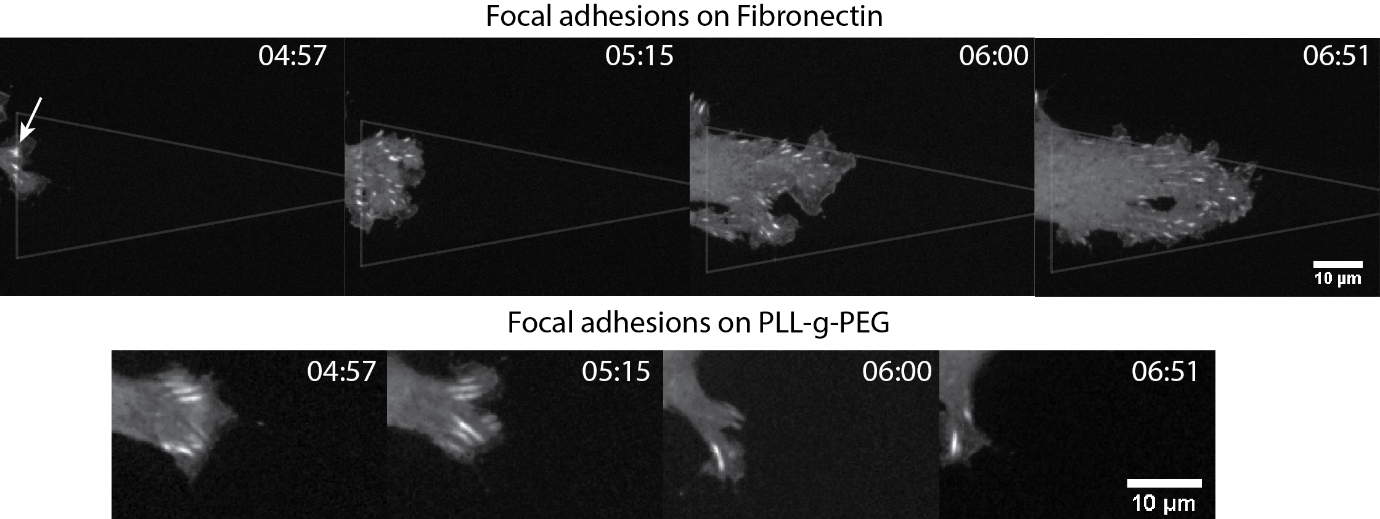

**Figure S3: Time-lapse sequences of focal adhesions on fibronectin triangles and on glass coated with PLL-g-PEG.** Protrusions come from the same cell at the same time points on both fibronectin triangles and on pLL-g-PEG. After a first docking line of focal contacts assembled on the edge of the triangle (white arrow), a wave of adhesion sites appears, promoting migration towards this direction. On PLL-g-PEG (below), new contacts cannot be nucleated beyond the first row of focal adhesions, and this leads to retraction of the protrusion.

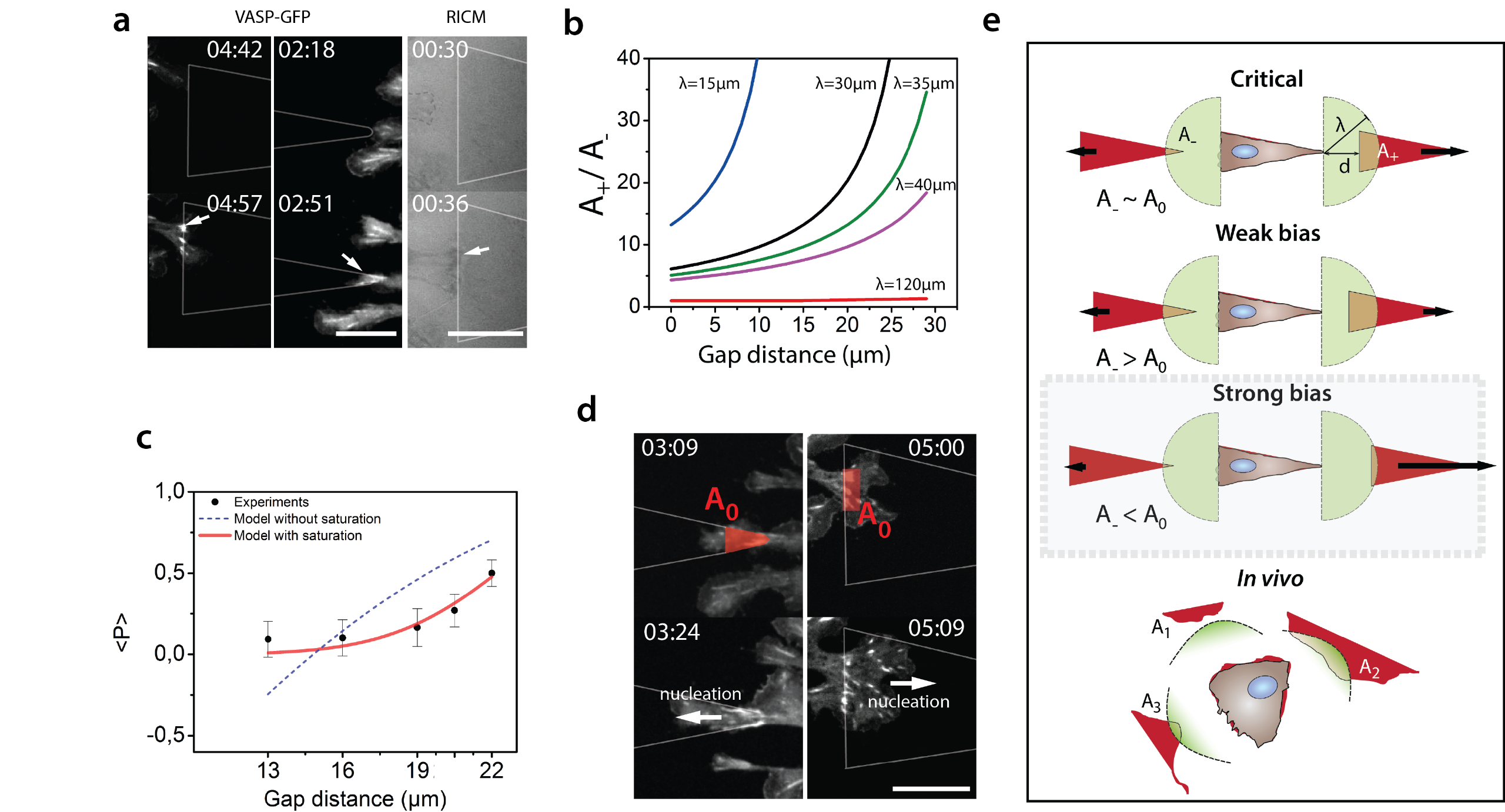

**Figure S4:** Theoretical ratio of probed areas as a function of gap distances for various protrusion lengths λ.
